# Multi-model biological and sequence information fusion for gene regulatory network inference from single-cell transcriptomics

**DOI:** 10.64898/2026.09.13.751326

**Authors:** Lin Zhong, Bin Yan, Junwen Wang, Minzhu Xie

## Abstract

Identification of transcription factor–target gene interactions and construction of the gene regulatory networks (GRNs) are essential for understanding the molecular mechanisms underlying transcriptional gene regulation. Large-scale single-cell transcriptomics across different tissues offers unprecedented resolution of cellular diversity and regulatory dynamics by capturing gene expression heterogeneity. However, existing methods often lack effective multimodal integration and fail to fully exploit the hierarchical structure in Gene Ontology (GO) and gene sequence level representations, which limits their ability for predictive performance and biological interpretability. We present scMGFGRN, a multi-model deep learning framework that integrates single-cell transcriptomic profiles with gene functional hierarchical relationships, gene sequences by leveraging denoising auto-encoders, graph attention feature extraction and pertained DNA language model to capture multi-source dependencies within multi-model biological knowledge, while its gated multi-head attention module effectively identifies informative regulatory signatures and integrate complementary features from different sources to predict accurate gene regulatory networks. Benchmarking on the seven datasets of human and mouse demonstrates that scMGFGRN out-performs state-of-the-art methods in identifying GRNs. Further analyses reveal that scMGFGRN effectively identifies novel TF– gene interactions (TGIs) and reconstructs cell-type-specific GRNs. Interpretability analysis reveals the contribution patterns of heterogeneous biological sources, demonstrating the ability of scMGFGRN to integrate transcriptomic profiles with multi-model structure information.

## I. INTRODUCTION

The architectural integrity of eukaryotic cellular function is governed by a multilayered regulatory complex, where transcription factors (TFs) and their target genes engage in a precise regulatory mechanism of activation and inhibition. These interactions collectively form complex gene regulatory networks (GRNs), which serve as the foundation of cellular regulation. GRN inference plays a critical role in functional gene annotation, cancer biomarker discovery, and target-driven drug development [1]. Therefore, deciphering the mechanistic principles of these TF–gene interrelationships and capturing the dynamics of GRNs are crucial for unraveling the regulatory mechanisms underlying biological processes and complex biological phenomena.

Numerous methods have been developed to predict GRNs from single-cell transcriptomic data to reveal transcriptional regulatory relationships. Single-cell network synthesis (SCNS) [2] applied a Boolean network model to reconstruct GRNs. SCODE [3] utilized ordinary differential equations to reconstruct GRNs, leveraging pseudotime as high-resolution temporal information. Tools such as PCC [4], PIDC [5], and PPCOR [6] employ single-cell transcriptomic data to model pairwise mutual information between genes, subsequently calculating the ratio of unique components to mutual information. Machine learning-based approaches, including GENIE3 [7], GRNBoost2 [8], and SCENIC [9], apply algorithms such as gradient boosting machines and random forests to treat TF– gene interaction prediction and GRN inference as large-scale regression or classification problems. These methods are computationally intensive, which limits their broader application.

Distinct from traditional methods, deep learning-based models can integrate not only gene expression profiles but also known regulator–gene interactions and organism/tissue or cell-type-specific information [10–12]. For instance, the ANN-based gene network embedding (GNE) [13] integrates expression data and the topological structure of gene interactions into latent representations to predict regulatory interactions. CNN-based co-expression analysis [14] integrates prior knowledge and other information, encoding gene expression data into matrices to extract spatial features for GRN inference. DeepRIG [15] employed a graph autoencoder to embed co-expression and regulatory information into latent representations and reconstruct GRNs. DeepSEM [11] reconstructs cell-type-specific GRNs autonomously using a beta-VAE, showcasing remarkable potential for handling complex and uncertain single-cell data. Recently, methodologies such as GENELink [16] have further harnessed the strengths of graph neural networks for GRN inference. scMGATGRN [17] utilizes a multi-view graph attention network for inferring GRNs from single-cell transcriptomic data. However, these models are highly sensitive to the quality of the input graph. Inaccurate prior knowledge or noisy initial co-expression data can undermine the robustness of the model.

To address these issues, we propose a novel deep learning framework named scMGFGRN, which integrates single-cell transcriptomic profiles with multi-model structured biological knowledge, including gene function hierarchical relationships and gene sequence information and dynamic pseudotimes series for GRN inference. The scMGFGRN architecture employs a denoising auto-encoder (DAE) to alleviate technical noise and extract gene expression features in single-cell transcriptomic data and a bidirectional gated recurrent unit (Bi-GRU) to capture complex time-series regulatory dynamics from pseudo-time-ordered expression profiles. Furthermore, graph-based feature extraction is introduced to model hierarchical structure within GO terms, enabling the learning of informative gene functional representations. A gated multi-head attention module is further incorporated to adaptively identify informative regulatory signatures from fused multi-model latent representations and improve the prediction of gene regulatory relationships. We evaluated the performance of scMGFGRN using cross-validation across different species and diverse cell-type datasets, comparing it with state-of-the-art methods and the results demonstrate that scMGFGRN consistently outperforms other methods in GRN inference. Overall, the main contributions of this paper are summarized as follows:

- We designed the scMGFGRN framwork, a novel multi-model fusion framework integrates single-cell transcriptomics with GO hierarchical relationships and gene sequences for accurate GRN inference and a gated multi-head attention mechanism captures representative regulatory features and adaptively combines multi-model biological information.
- We apply a denoising auto-encoder for noise reduction and feature extraction of gene expression patterns from single-cell transcriptomic data. To encoder the hierarchical relationships of the gene function, we designed the multi-hot encoding module to capture complex structure representations of gene function and employ a graph attention module to enhance graph-based feature extraction. Additionally, a DNABERT language model is employed to capture gene sequence feature for GRN prediction.
- Comprehensive experiments on multiple scRNA-seq datasets demonstrate that our method outperforms the state-of-the-art methods in GRN inference. The discovery of cell-type-specific networks and unknown potential TF-target relationships provides deeper biological insights into gene regulatory mechanisms.

## II. DATASET

To evaluate the performance of scMGFGRN, we utilized seven single-cell RNA sequencing (scRNA-seq) datasets widely adopted for benchmarking deep learning algorithms [16, 18]. These include: (i) human embryonic stem cells (hESC); (ii) human mature hepatocytes (hHep); (iii) mouse dendritic cells (mDC); (iv) mouse embryonic stem cells (mESC); (v) mouse erythroid hematopoietic stem cells (mHSC-E); (vi) mouse myeloid hematopoietic stem cells (mHSC-GM); and (vii) mouse lymphoid hematopoietic stem cells (mHSC-L).

Each of these seven datasets incorporates three types of ground-truth networks from functional interaction networks that were sourced from the Search Tool for Recurring Instances of Neighbouring Genes (STRING) [19], nonspecific ChIP-seq [20–22] and cell-type-specific ChIP-seq [23–25]. Additionally, the mESC dataset includes a functional loss-of-function/gain-of-function (LOF/GOF) ground-truth network [25]. BEELINE is currently the most widely applied data preprocessing framework [18]. We employed BEELINE to preprocess each scRNA-seq dataset and infer interactions originating from TFs. We filtered the top 500 and 1,000 genes with the most significant expression variations for model performance evaluation. To incorporate structured regulatory information and priors into our model, we integrated the gene promoter sequences and hierarchical structure of the GO terms including the Gene Ontology Biological Process (GO:BP), Molecular Function (GO:MF) and Cellular Component (GO:CC) [26].

Feature representation is essential for model training. We compute the PCC for the transcriptomic expression profile. For scRNA-seq, the feature of each gene (including TFs) corresponds to a set of its expressions in all samples. To acquire the information containing internal correlations, we calculated PCCs of mRNA expression between TFs and genes, and the expression similarity matrix were generated. The PCC of a pair of genes/TFs expression i and j is calculated as follows:

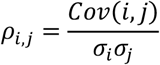

where Cov(*i, j*) denotes the covariance between gene/TF *i* and gene *j*, and *σ*_*i*_, *σ*_*j*_ are their respective expression standard deviations. Each row of the resulting similarity matrix captures the regulatory affinity between a given TF and all other genes and serves as the input for downstream feature extraction. For the pseudotime transcriptomic data, we calculated the pseudotime ordering from each expression profiles and subsequently arranged the cells according to this order to capture temporal time-series feature from the transcriptomic data. We applied SlingShot [27] to calculate pseudotimes of each dataset. The pseudotimes are time-series information of ‘each cell, which reflect the relative order in the whole differentiation process.

## III. METHOD

As shown in Fig.1, the framework integrates single-cell RNA-seq profiles, GO hierarchical information, gene sequence knowledge, and pseudotime expression patterns to infer gene regulatory networks. Transcriptomic features are denoised and compressed using a denoising auto-encoder, while temporal regulatory patterns from pseudotime-ordered expression profiles are captured by a bidirectional gated recurrent unit. A graph-based GAT module encodes GO hierarchy leveraging the directed acyclic graph (DAG) to produce robust hierarchical biological representations, whereas gene sequences are encoded using the pretrained DNABERT-2 model. The heterogeneous latent features are subsequently integrated by a gated multi-head attention module to identify informative regulatory signatures, and a fully connected layer predicts TF-target gene relationships.

### A. Denoising auto-encoder with attention mechanism

Denoising auto-encoder (DAE) [28] is an unsupervised learning algorithm, the basic principle of which is to achieve data noise reduction and feature extraction by introducing noise into the input data and trying to reconstruct the original, noise-free data from the noisy data [29]. To extract gene regulatory signatures from noisy, high-dimensional transcriptomic data, we designed a denoising autoencoder architecture with attention mechanism, as illustrated in Fig. 1B. Unlike conventional DAEs, our model integrates multi-head attention (MHA) [30] and gated linear units (GLU) [31] to enhance the capture of multi-source TF-gene dependencies. The DAE effectively eliminates redundancies and dropout noises within scRNA-seq data, providing a denoised latent representation of gene expression for downstream GRN inference. We introduced dropout noise by randomly setting a fraction of the input *x* elements to zero, as defined below.

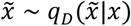

**Figure 1.**
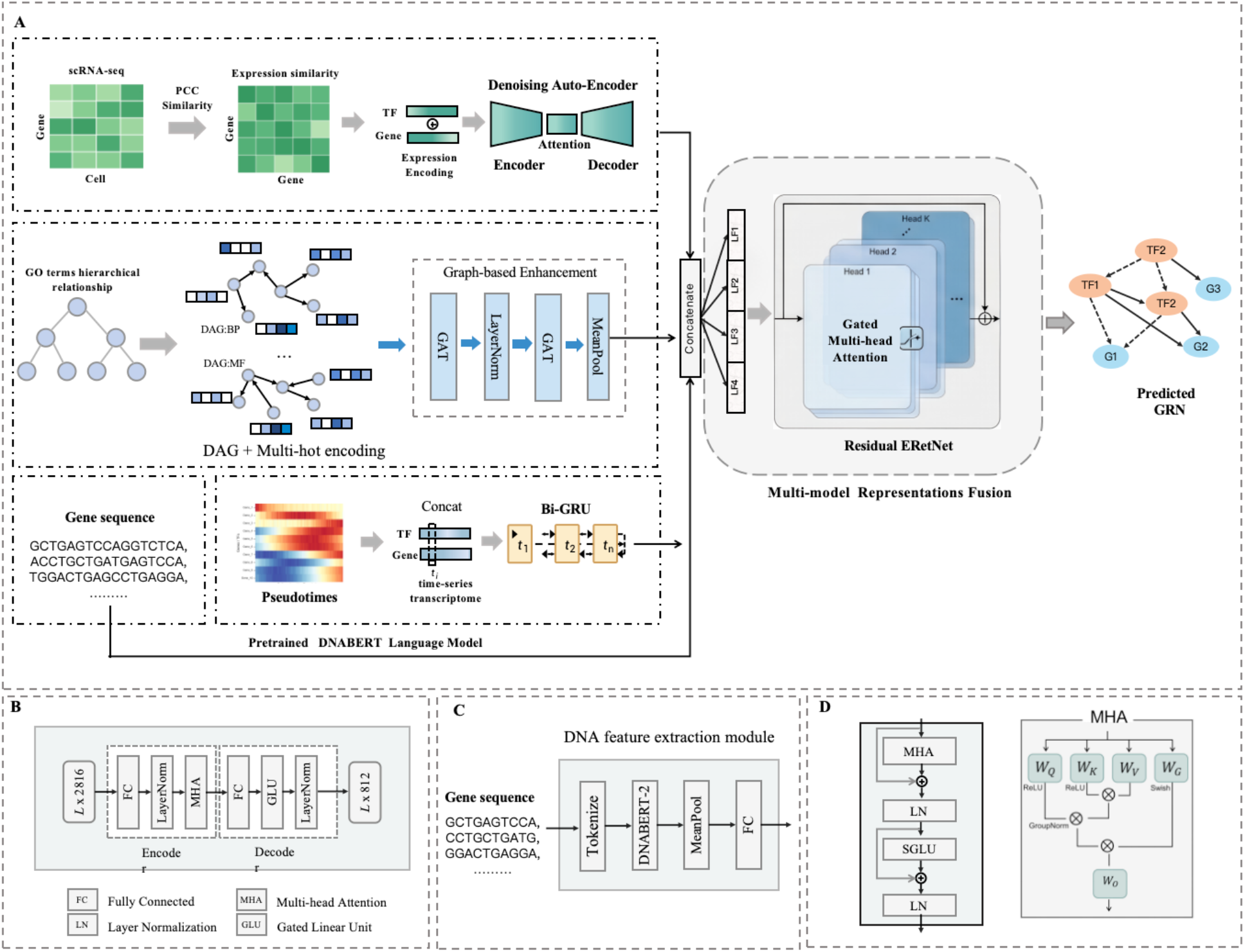
The architecture overview of scMGFGRN **(A)** This framework employs multi-modal deep learning strategy that integrates single-cell transcriptomics, ontological hierarchies, promoter sequence knowledge, and pseudotime expression dynamics for genome-scale regulatory network inference. By leveraging dedicated neural encoders to extract multi-model representations and a gated attention-based fusion mechanism, the model effectively captures heterogeneous biological signals to predict transcription factor–target gene interactions with high accuracy. **(B)**, Architecture of the denoising auto-encoder module. The decode and encoder architecture learns robust transcriptomic representations by reconstructing inputs and reducing the influence of noise in sparse single-cell expression profiles. **(C)**, Architecture of the DNABERRT-2 encoding module. **(D)**, Architecture of the residual gated attention fusion module. Multi-model latent representations are integrated through gated multi-head attention, enabling the model to extract regulatory-relevant features from heterogeneous biological information for GRN prediction.

The encoder aims to project the corrupted input 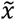 into a lower-dimensional latent space. We add a MHA module to enhance feature extraction.

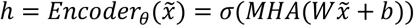

The decoder then attempts to reconstruct the original noise-free input *x* from the hidden representation *h*:

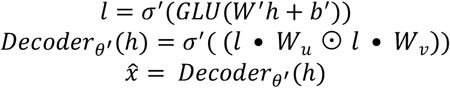

The module is trained by minimizing the Mean Squared Error Loss, which quantifies the discrepancy between the original input and its reconstruction:

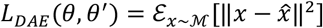

### B. Gene functional hierarchical relationships extraction

Gene Ontology provides a comprehensive and hierarchically organized knowledge framework for describing gene functions, including biological processes (BP), molecular functions (MF), and cellular components (CC). GO terms are structured through complex semantic relationships, forming a directed acyclic graph (DAG). To incorporate hierarchical knowledge into GRN inference, we developed an ontology-aware representation encoding module that integrates structural information of GO terms. Specifically, the graph is constructed as *G* = (*V, E*), where each node represents a GO term and edges denote “is_a” hierarchical relationships between terms. For each gene-associated term *g*_*i*_, a multi-hot structural vector is generated by encoding its ancestor terms and itself in the hierarchy. This structural representation captures the hierarchical context of gene functions and is further projected into a latent feature space through a linear transformation. The construction of multi-hot vector and structural embedding are as follows:

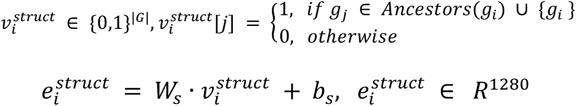

where G denotes the full set of GO terms, *g*_*j*_ represents a specific GO term, and Ancestors (*g*_*j*_) denotes all ancestor terms of *g*_*j*_ in the the GO hierarchy. 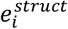 is a low-dimensional, dense structural embedding, 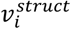 is a multi-hot sparse vector that denotes the ancestor structure of 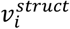, and *W*_*s*_, *b*_*s*_ are the learnable weight matrix and bias.

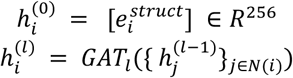

Here, 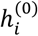 denotes the initial feature of term *i*, while 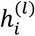 corresponds to its representation after the *l*-th GAT layer, and *N*(*i*) is the set of its neighboring nodes. Through iterative message passing along the GO DAG, each term progressively aggregates structural representations from both its parent and child terms, enabling the encoder to capture hierarchical information beyond independent term-wise embeddings. A LayerNorm operation and global average pooling over all nodes is then applied to produce the final GO graph-level embedding for TF and gene representation enhancement. The detailed procedure is as follows:

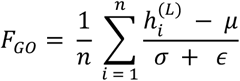

where *n* is the number of GO terms, 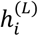 is the final GAT embedding of term *i*, and *μ, σ*^2^, and *ϵ* are the mean, variance, and a small constant of the input features.

### C. Bi-GRU module for time-series feature extraction

To capture the dynamic pseudo time-series dependencies within transcriptomic profiles, we employed a bidirectional gated recurrent unit (Bi-GRU) [32]. Unlike standard GRUs that only process historical data, the Bi-GRU architecture integrates both preceding and succeeding contextual information, providing a comprehensive representation of pseudotime gene expression correlation vectors. The formula for bidirectional contextual information extraction is defined as follows:

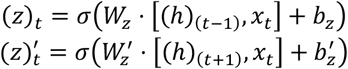

where *x*_*t*_ is the current input vector, (*h*)_(*t*−1)_, (*h*)_(*t*+1)_ are the succeeding unit state at time-step *t* − 1 and the preceding unit state at time-step *t* + 1. *σ* denotes the sigmoid activation function.

The final hidden state 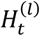 at time step *t* is obtained by concatenating the forward hidden state 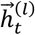 and the backward hidden state 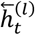, [a,b] denotes the concatenation operation:

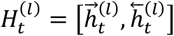

Specifically, we implemented a Bi-GRU architecture to capture multi-level hierarchical time-series features and complex temporal dependencies from the correlation vectors. The Bi-GRU layer takes the ordered TF expression and target expression as input, while each subsequent layer receives the concatenated hidden states from the previous layer as its input. Finally, the concatenated hidden representations from all time steps are passed through a mean pooling layer and a fully connected layer to derive the time-step features.

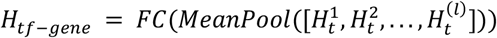

### D. Gene sequence feature extraction using DNABERT-2

Gene sequences contain abundant regulatory information, especially in promoter area, in which sequences are enriched with diverse regulatory signals, including transcription factor binding sites (TFBS), cis-regulatory elements, and conserved motifs, which collectively encode critical information for gene regulation [33]. To incorporate regulatory sequence information into scMGF-GRN, we extracted promoter sequences for each target gene based on the transcription start site (TSS). For each gene, the promoter region was defined as the genomic interval spanning 1,500 bp upstream and 500 bp downstream of the TSS. These promoter sequences were then encoded using DNABERT-2 [34] to capture sequence-level regulatory features. Finally, the promoter-derived embedding was projected into the model’s feature space and integrated through the gated multi-head attention module. The procedure for sequence feature extraction using the pre-trained DNA language model DNABERT-2 is defined as follows:

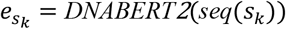

where seq(*S*_*k*_) denotes the gene promoter sequence.

### E. Gated multi-head attention mechanism for feature fusion

The multi-head attention mechanism [30] is an extension of self-attention that enables the model to jointly attend to information from different representation spaces. However, standard MHA may struggle with the inherent noise and high dimensionality of single-cell transcriptomic data. To address this, we employ a gated multi-head attention (Gated MHA) module [35]. The gated variant incorporates a gating mechanism to dynamically regulate the flow of information and acts as a filter that assigns higher learned weights to critical gene regulatory features while suppressing noise and less relevant dependencies, which can lead to an improved model performance and a smoother training process. The formulas are defined as follows.

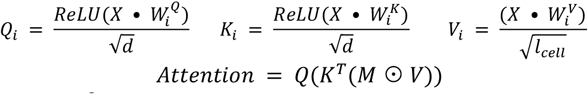

where 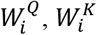 and 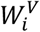 denote the weight matrices, *l*_*cell*_ is the number of the cell, *M* is the fused TF-gene representation. To stabilize deep training and refine latent feature decomposition, we evolved the ERetNet architecture by replacing the standard pre-layer normalization with DeepNorm [36]. This transition, coupled with our Gated MHA module, enables the model to effectively prioritize pivotal TGIs while maintaining numerical stability during optimization.

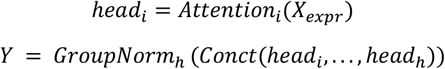

In addition, we add a swish gate [37] *W*_*G*_ to increase the non-linearity of ERetNet. where ⊙ is the element-wise product.

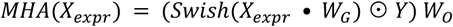

Finally, the local auto-encoder features *LF*_1_ and time-series features *LF*_2_ of the TF and gene are fused with GO embedding *LF*_3_ and promoter sequence embedding *LF*_4_ obtained from the above steps are fused in the Gated MHA module, based on the formulas:

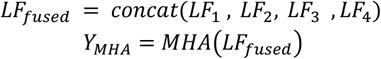

To enhance representation learning and stabilize training dynamics, we evolved the standard feed-forward network in ERetNet into a gated bilinear framework. By incorporating a Gated Linear Unit (GLU) [31], the model gains a multiplicative gating mechanism that modulates feature propagation and facilitates superior channel integration.

Furthermore, recognizing that the gating interaction itself provides sufficient non-linear inductive bias, we adopted the Simplified GLU (SGLU), which omits the Swish activation.

This architectural refinement significantly reduces computational overhead and accelerates throughput without compromising modeling expressive power, achieving an optimal balance between performance and efficiency. The SGLU is defined as follows:

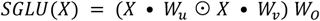

The residual module can partially prevent the problem of gradient disappearance in neural networks and enables scMGFGRN’s network to be designed deeper. We apply the residual adding on the Gated MHA module and the new post-norm normalization method DeepNorm [38]. DeepNorm reduces the contribution ratio of each network block to the output, thereby reducing the amount of gradient that needs to be updated and ensuring the stability of training.

We infer GRNs as a prediction task. Each TF–gene pair corresponds to two results, interact or not interact, so the prediction task is a binary classification. The datasets used in the analysis are all balanced. The binary cross entropy loss is applied to train the model, and the formulas are listed as follows:

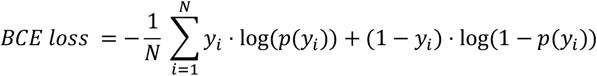

where y_i_ and p_i_ represent the label and prediction corresponding to the ith data, N is the number of the training sample. Besides, Mean Squared Error Loss is also employed to be an auxiliary loss which regularizes the extra output of the denoising auto-encoder.

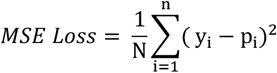

### F. Model training and performance evaluation

To ensure a robust and unbiased evaluation, we implemented a balanced sampling strategy across diverse cell lineages and species. Positive samples were derived from gold-standard regulatory networks, while an equal number of negative samples were randomly generated. To mitigate the impact of potential false negatives, we employed 5-fold cross-validation, partitioning the dataset into training and testing sets at a 4:1 ratio. The implementation environment and hyper-parameter configurations of the model are detailed in Supplementary Table S3. We use the Area Under the Receiver Operating Characteristic (AUROC) and Area Under the Precision-Recall Curve (AUPRC) to evaluate model performance. True Positive Rate (TPR), False Positive Rate (FPR), the Precision and Recall are defined by equations:We compared scMGFGRN with the PCC [4], GENIE3 [7], GRNBoost2 [8], SCENIC [9], SCODE [3], DeepSEM [11], GNE [13], DeepRIG [15], GENELink [16] and scMGATGRN [17] to test the performance of scMGFGRN. These methods include shallow learning algorithms (PCC, GENIE3, GRNBoost2, SCENIC, SCODE) and state-of-the-art deep learning-based methods (DeepSEM, GNE, DeepRIG, GENELink, scM-GAGRN).

## IV RESULT

### A. Comparison with state-of-the-art methods

We conducted experiments on the integration of single-cell expression profiles, GO:BP hierarchy, and sequence data first. As shown in Fig. 2, scMGFGRN outperforms all competing methods in terms of the average AUROC across all datasets. Specifically, on the STRING (TFs + 500) dataset, scMGFGRN achieves an average AUROC of 94.03%, surpassing the second-best method scMGATGRN (91.19%) by 2.84%. On the STRING (TFs + 1000) dataset, scMGFGRN obtains 93.18%, outperforming scMGATGRN (90.93%) by 2.25%. The Area Under the Receiver Operating Characteristic curve (AUROC) and Area Under the Precision-Recall Curve (AUPRC) on mESC dataset were shown in Fig. 3A and Fig. 3B. scMGFGRN consistently outperformed all competing methods, achieving the highest AUROC score of 0.9456. On the nonspecific ChIP-seq (TFs + 500) dataset, scMGFGRN reaches 88.36%, exceeding scMGATGRN (81.28%) by 7.08%. On the nonspecific ChIP-seq (TFs + 1000) dataset, scMGFGRN achieves 89.81%, outperforming the second-best method GENELink (83.98%) by 5.83%.

**Figure 2.**
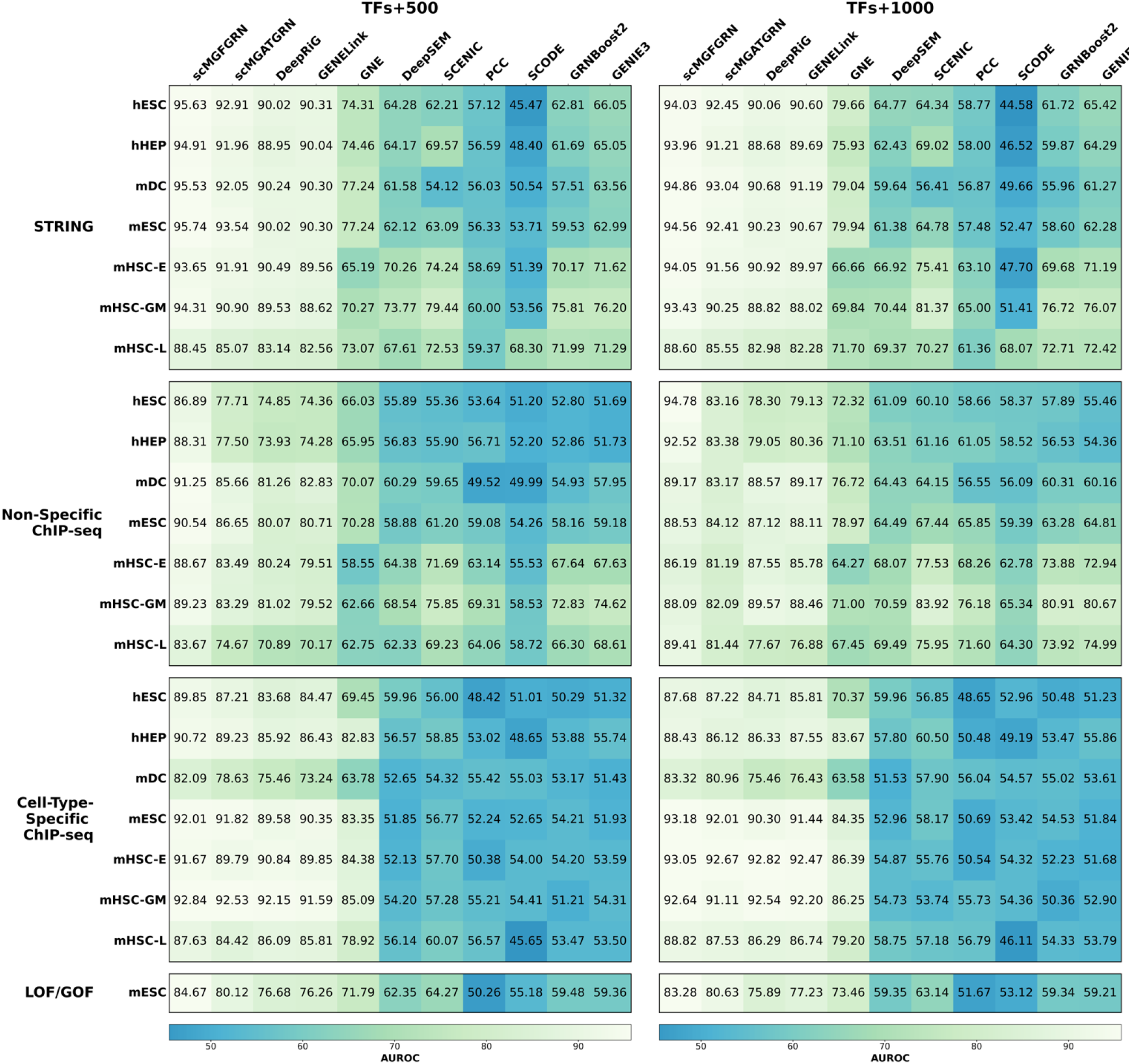
Performance comparison across diverse datasets and ground-truth standards. The heatmaps present the AUROC scores for scMGFGRN and eleven baseline methods across seven benchmark datasets, evaluated against four types of ground-truth networks: STRING, nonspecific ChIP-seq, cell-type-specific ChIP-seq, and LOF/GOF.

**Figure 3.**
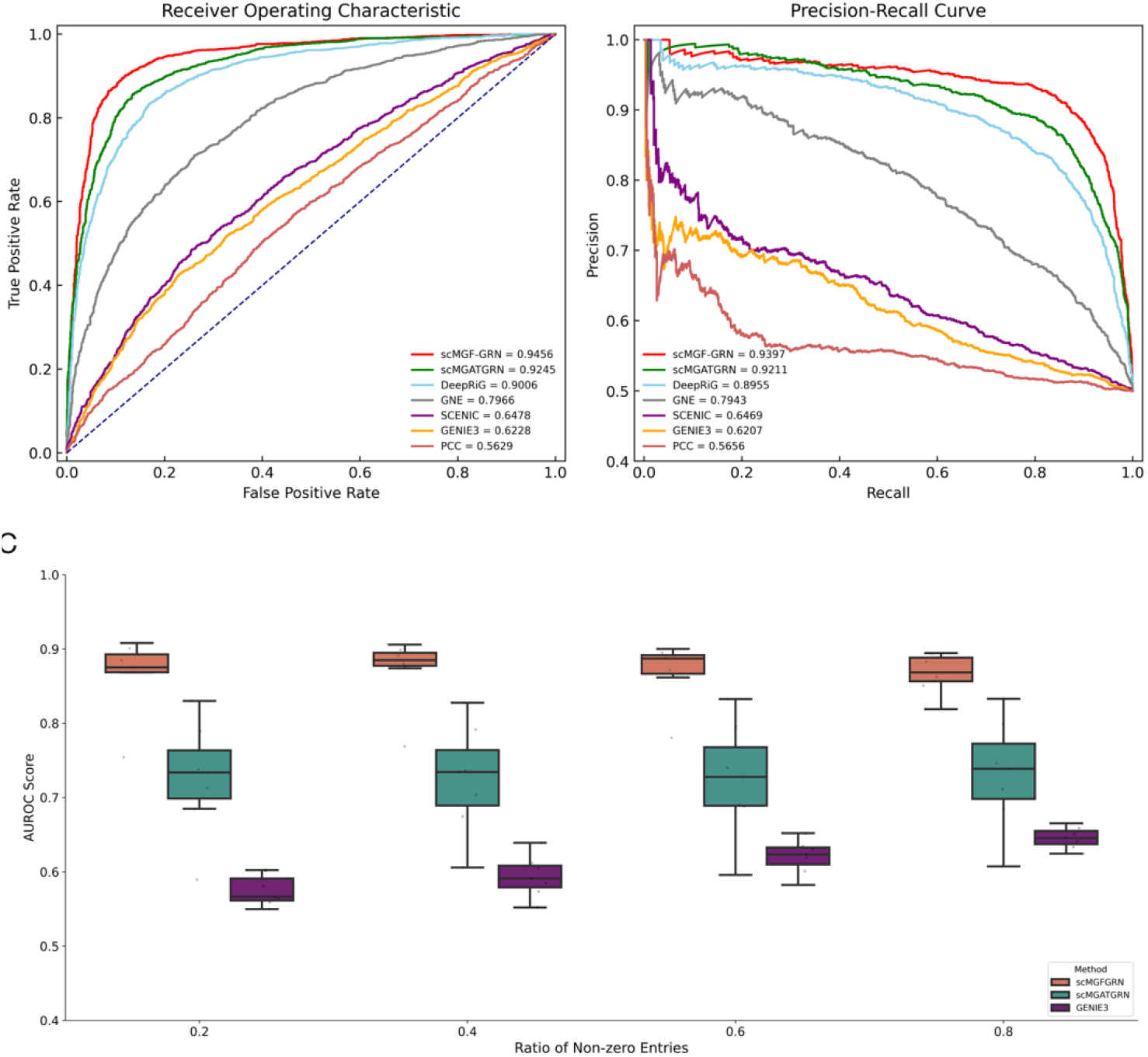
(A),(B), ROC and P-R curves comparison of scMGFGRN against six baseline methods across benchmark datasets. scMGFGRN outperforms methods like scMGATGRN and DeepRiG in GRN inference. **(C)**, Sparsity test for evaluating model robustness across varying data densities. The boxplots illustrate the AUROC scores of scMGFGRN, scMGATGRN and GENIE3 across seven benchmarking datasets under four different data density levels. We randomly set a specific proportion of the gene expression data to zero, resulting in four density levels. The ratio of non-zero entries in the gene expression matrices was controlled at 0.2, 0.4, 0.6, and 0.8

For the second-part ground-truth datasets (cell-type-specific ChIP-seq and LOF/GOF), as shown in Fig. 2, scMGFGRN out-performs all competing methods in terms of average AUROC across all cell-type-specific datasets. On the cell-type-specific ChIP-seq (TFs + 500) dataset, scMGFGRN achieves an average AUROC of 89.54%, surpassing the second-best method scMGATGRN (87.66%) by 1.88%. On the cell-type-specific ChIP-seq (TFs + 1000) dataset, scMGFGRN obtains 89.59%, outperforming scMGATGRN (88.23%) by 1.36%. The LOF/GOF datasets exhibit broadly similar predictive performance across the evaluated methods, achieving the highest AU-ROC of 84.67% and 83.28% respectively. As illustrated in Fig. 1, scMGFGRN consistently outperforms supervised learning methods and unsupervised methods in terms of AUROC on all cell-type-specific ChIP-seq and LOF/GOF datasets.

### B. Robustness analysis under varying data sparsity

To evaluate the resilience of scMGFGRN against the inherent technical noise and high dropout rates in single-cell transcriptomics, we conducted a systematic robustness stress test on seven single-cell benchmarking. We compared our model with two representative categories of GRN inference methods: scMGATGRN and GENIE3. We simulated four levels of data density, defined by the ratio of non-zero entries, ranging from 0.2 to 0.8. To further challenge the models’ denoising capabilities, we utilized nonspecific datasets characterized by high technical variance and non-biological signals. Performance was quantified by AUROC score across seven benchmark datasets.

The results were summarized in Figure 5C, across all sparsity levels, scMGFGRN consistently achieved the highest median AUROC (avg. = 0.87). Even in the most extreme scenario, our model maintained exceptional performance, whereas competing methods scMGATGRN and GENIE3 showed significant degradation. This suggests that the integrated DAE successfully captures the underlying regulatory mechanism despite severe information loss. In contrast, scMGATGRN showed significantly elongated boxes and whiskers. This contrast demonstrates that while traditional graph-based models like scMGATGRN struggle with sparsity and drop noise of scRNA-seq data, our DAE-integrated architecture effectively filters out redundant signals, ensuring robustness in complex regulatory environments. In conclusion, these results demonstrate that scMGFGRN is not only more accurate but also more robust than current state-of-the-art methods, making it a more reliable tool for analyzing noisy and sparse single-cell atlases.

### C. Analysis of architectural hyperparameters setting

To investigate the influence of the model architecture on GRN prediction performance, we evaluated three important architectural hyperparameters, including the GAT embedding dimension, number of attention heads, and number of attention network layers. These parameters directly determine the representation capacity, feature representation capability, and architectural depth of the proposed gated attention-based fusion module. For each parameter, the remaining model settings were kept unchanged, and the performance was evaluated using AU-ROC, ACC, and AUPR the hESC-500 benchmark datasets using ground-truth networks STRING, Non-specific and Specific networks.

We first evaluated embedding dimensions of 128, 256, and 512. As shown in Fig. 4A, increasing the dimension from 128 to 256 improved performance, whereas further increasing it to 512 resulted in slight declines in most settings. This suggests that a larger feature space initially improves the representation of heterogeneous biological information, while excessive dimensionality may introduce redundant features and increase model complexity. Therefore, 256 was selected as the embedding dimension. Next, we evaluated 4, 6, and 8 attention heads. As illustrated in Fig.4B, increasing the number of heads from 4 to 6 improved the overall performance, with the best AUROC, ACC, and AUPR increasing from 0.942, 0.895, and 0.898 to 0.948, 0.907, and 0.915, respectively. Finally, we tested 2, 3, 4, and 5 attention layers. As shown in Fig. 4C, performance generally improved with increasing depth up to 4 layers. For example, AUROC increased from 0.922 to 0.947, while adding a fifth layer led to a slight decline. This indicates that moderate network depth provides sufficient capacity to model higher-order interactions, whereas excessive depth may introduce redundant transformations and hinder optimization.

**Figure 4.**
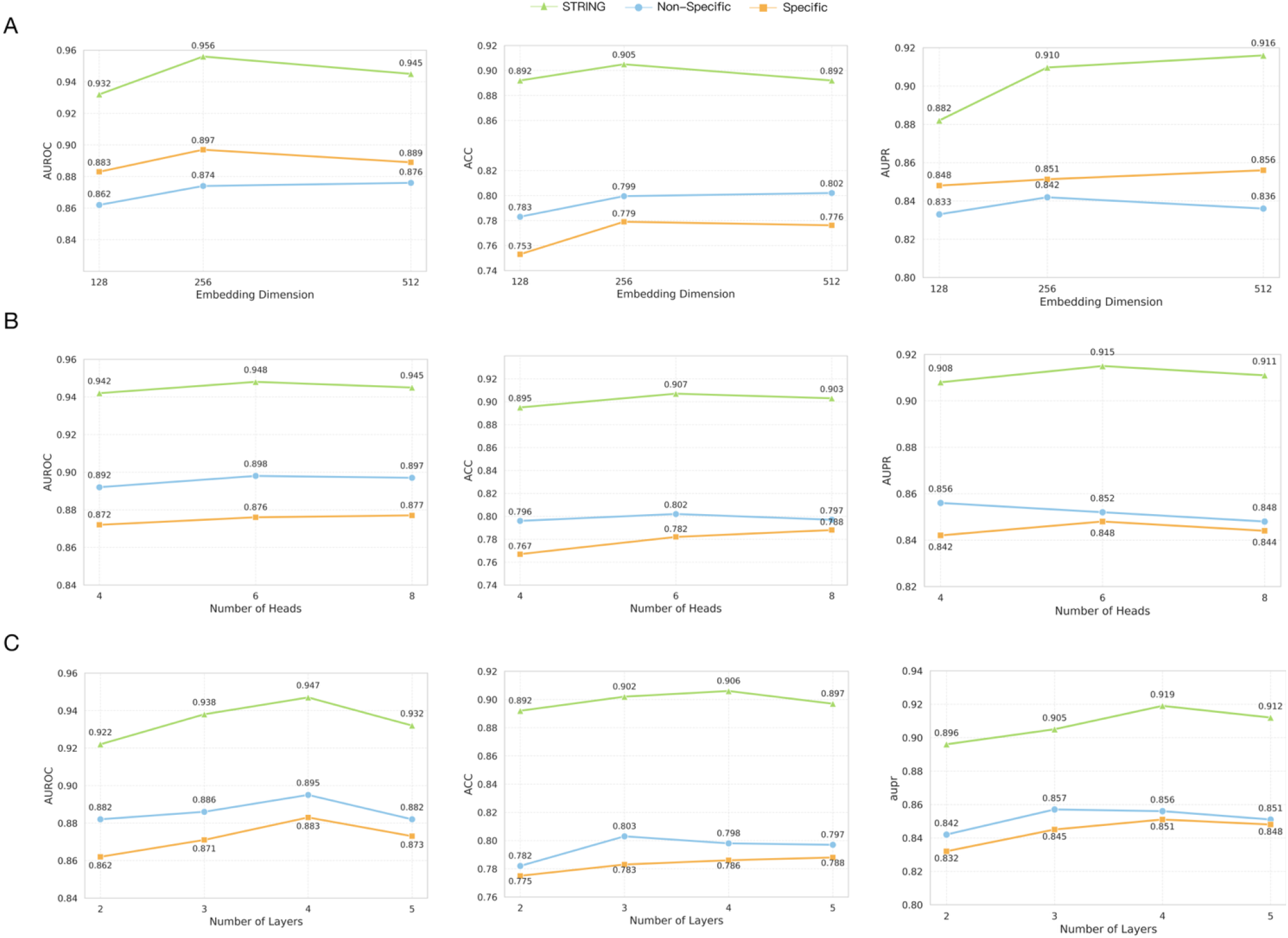
(A), Performance of the model on the hESC-500 dataset with different embedding dimension, varying number of attention heads and varying numbers of gated attention fusion modules, respectively, measured by AUROC, ACC, and AUPR scores under each configuration.

Overall, the hyperparameter analysis demonstrates that the proposed architecture benefits from a moderate representation dimension, a sufficient number of parallel attention heads, and an intermediate network depth. Based on the consistently favorable performance across AUROC, ACC, and AUPR, we selected an embedding dimension of 256, 6 attention heads, and 4 layers as the default configuration. This configuration provides sufficient capacity to capture complex interactions among heterogeneous biological features while avoiding the additional redundancy and complexity associated with excessively large or deep architectures. To further validate the generalization ability of the model, we then extended our feature fusion strategy to the other two primary GO domains: Molecular Function (GO:MF) and Cellular Component (GO:CC). As depicted in Figure 5, incorporating either GO:MF or GO:CC hierarchy alongside single-cell expressions yielded substantial performance enhancements over the baseline model, with AUROC scores consistently exceeding 0.90 across most cellular datasets. The integration of cellular component hierarchy demonstrated a marginally stronger predictive advantage compared to molecular functions in the majority of contexts. Most notably, when synchronously combined with gene promoter sequences, both the MF-driven and CC-driven joint frameworks converged on peak inference accuracy, frequently improving the AUROC boundaries beyond 0.94. These extended empirical findings firmly validate that the predictive robustness of our framework is not domain-restricted; rather, it effectively synthesizes diverse hierarchical layers of GO terms, structural sequence to optimize GRN reconstruction.

**Figure 5.**
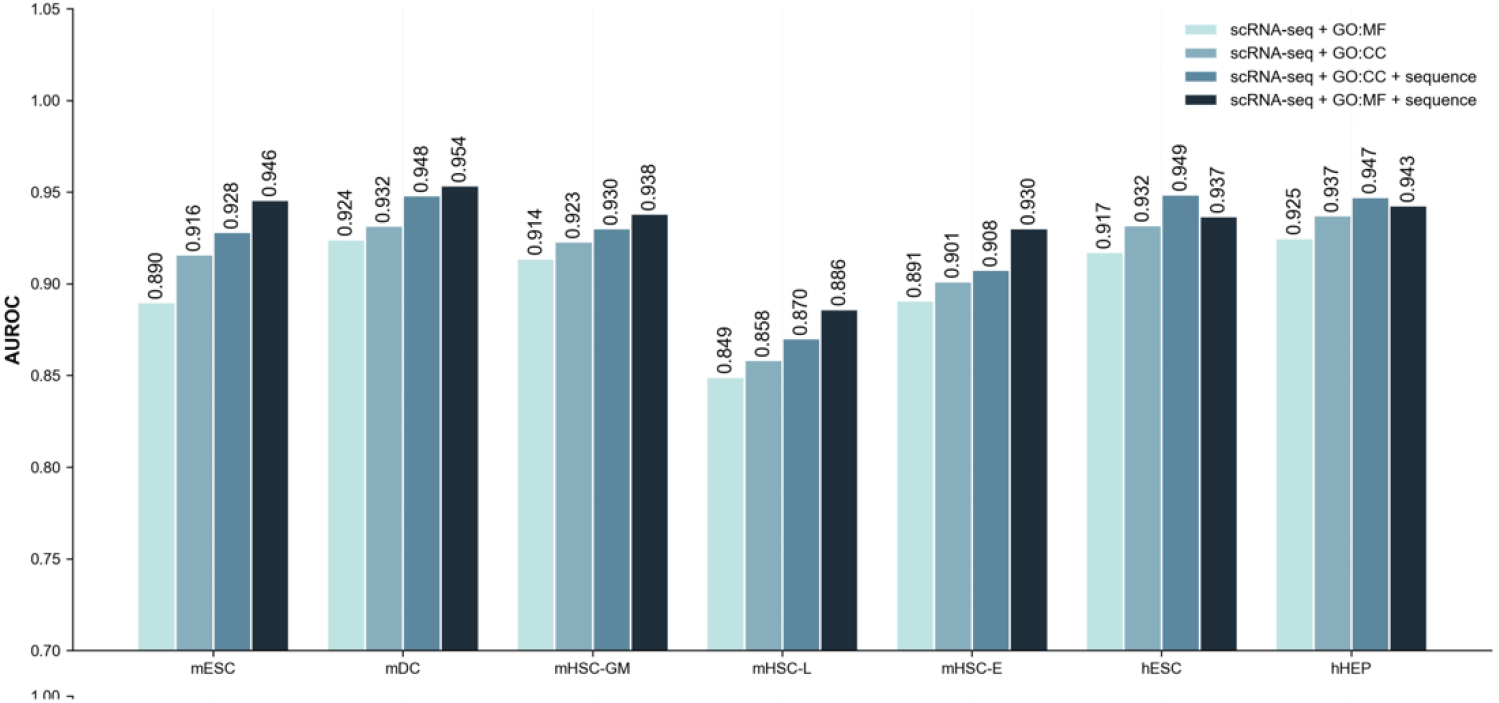
AUROC comparison of different GO domain integrations. Performance of scMGF-GRNS on seven datasets using singlecell data with GO:MF or GO:CC hierarchy, and in combination with promoter sequences.

### D. Ablation study of model components

To investigate the contribution of each component in scMGFGRN, we performed ablation experiments by progressively integrating different modules. As shown in Table 1, Bi-GRU alone achieved an AUROC of 0.8891, demonstrating that pseudotime-based sequential features provide valuable information for capturing dynamic regulatory patterns. Incorporating the DAE module further improved the performance of an AUROC to 0.9116, indicating that robust feature representations learned from noisy single-cell data enhance GRN inference. Integration of the DNABert module yielded an AUROC of 0.9248, while the GAT module further improved the combined model performance to AUROC 0.9397 when integrated with Bi-GRU and DAE, indicating that incorporating semantic and structure knowledge from GO can provide additional information for identification of regulatory interactions. Finally, the complete scMGFGRN model achieved the best performance, with AUROC, ACC values of 0.9486, 0.8813, respectively. These results demonstrate that each component contributes complementary information, and the multi-source fusion strategy effectively integrates transcriptional dynamics, GO hierarchy and sequence feature for accurate GRN inference.

**Table 1.**
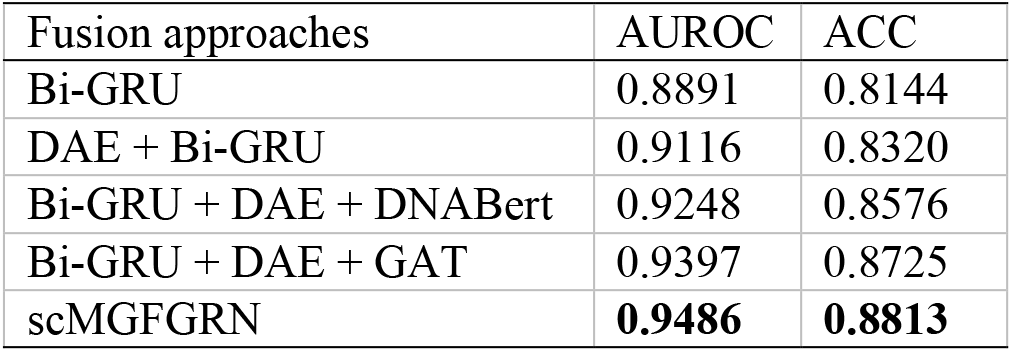
Ablation study of model components.

| Fusion approaches | AUROC | ACC |
| --- | --- | --- |
| Bi-GRU | 0.8891 | 0.8144 |
| DAE + Bi-GRU | 0.9116 | 0.8320 |
| Bi-GRU + DAE + DNABert | 0.9248 | 0.8576 |
| Bi-GRU + DAE + GAT | 0.9397 | 0.8725 |
| scMGFGRN | **0.9486** | **0.8813** |

### E. Cell-type specific gene regulatory network inference

To further explore the regulatory heterogeneity across different immune cell populations, we applied scMGFGRN to infer cell-type specific GRNs (ctGRNs). The scRNA-seq data were derived from human peripheral blood mononuclear cells (PBMCs), comprising over 8000 single cells from a healthy donor. Building upon previous research, we annotated cell types using marker genes reported in the literature [15]. The annotation identified four primary cell types: B cells, CD4 T cells, CD8 T cells and CD14 Monocytes. For establishing the prior networks of these cell types, we adopted regulatory relationships from the hTFtarget database, which integrates information from ChIP-seq, TF binding sites, and epigenetic modifications [39]. The predicted scores of TF-target pairs that are higher than 0.5 was considered for further investigation.

Core transcription factors are pivotal in cellular development and differentiation. By constructing ctGRNs for the three major cell types, we leveraged the out-degree of each gene to identify core TFs and their respective target genes for each cell type. The t-SNE visualization (Fig. 7A) of the PBMC scRNA-seq dataset reveals four distinct clusters corresponding to CD4 T cells, CD8 T cells and CD14 monocytes, demonstrating clear transcriptomic separation among the majorimmune cell types. We then infer the ctGRNs from the four specific single-cell clusters using scMGFGRN. As shown in Fig. 7B, the B cell specific network exhibits a highly centralized and interconnected core consisting of PAX5, BACH2, RUNX3, and SPIB. This central position in our ctGRN corroborates its known function of PAX5 in B-cell development and its implications in leukemogenesis [40]. The CD8+ T cell specific network (Fig. 7C) is characterized by a robust regulatory module centered around MYB, MEF2C, and BCL11A, highlighting its pivotal role in regulating CD8+ T cell stemness, survival, and longevity, which are critical for sustained antitumor immunity [41] and the maintenance of T-cell identity and development [42]. In the CD4+ T cell network (Fig. 7D), BACH2, RUNX3, and MYB emerged as the primary regulatory hubs, aligning with its established role as a critical hub TF for maintaining immune [43] and T-cell lineage development [44], reinforcing the model’s ability to capture essential developmental regulators.

**Figure 7.**
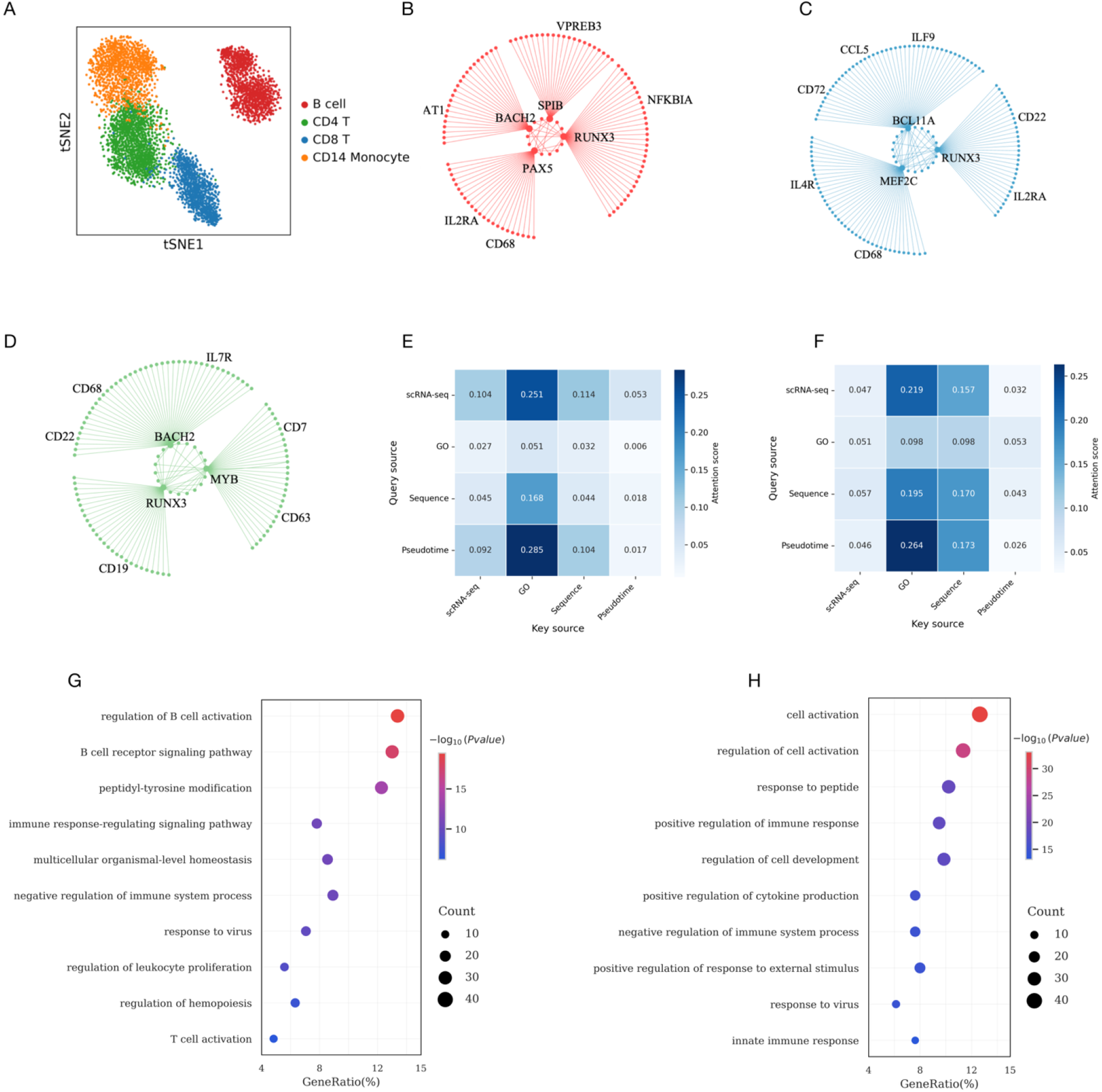
Application of scMGFGRN on PBMC scRNA-seq data for ctGRNs inference. (A), tSNE visualization of four cell types within the PBMC scRNA-seq dataset. (B),(C),(D), The ctGRNs of B cells, CD8 T cells and CD4 T cells and their core TFs and target genes identified by scMGFGRN. (E),(F), Heatmaps of attention score distributions for the two TF-gene pairs KLF6-JUN and MCM7-RIF1. (G),(H) Functional enrichment analysis of scMGF-GRNS inferred B cell regulatory networks and CD4 + T cells regulatory networks.

To delineate the specific functional profiles of adaptive immune subpopulations, GO biological process enrichment analyses were conducted for B cells and CD4+ T cells. As shown in Fig. 7G, distinct functional specialization was observed between the two lineages. B cells displayed higher significance in B cell receptor activation mechanisms, including regulation of B cell activation, B cell receptor signaling pathway, and peptidyl-tyrosine modification, which aligns with the fundamental requirement of B cells to translate extracellular antigen binding into intracellular. Furthermore, the enrichment of antigen processing and presentation of peptide antigen together with T cell costimulation highlights the specialized capacity of these B cells to act as professional antigen-presenting cells that modulate helper T cell fate. Conversely, CD4+ T cells showed preferential enrichment in cell activation processes, responses to peptide and viral, and regulations-related immune responses, which aligns with the fundamental requirement of CD4+ T cell to organize cell-mediated adaptive immune responses. These results based on inferred ctGRNs in B cells and CD4^+^ T cells demonstrate that scMGFGRN successfully integrates functional hierarchy to predict accurate dynamic GRNs. Moreover, the predicted GRNs exhibit strong functional consistency, as reflected by the concordance between the inferred regulatory relationships and the established GO biological process enrichments specific to each cell type, thus providing a reliable basis for deciphering cell-type-specific regulatory logic across diverse biological contexts.

### F. Interpretability analysis

To investigate the contribution and interaction patterns of heterogeneous biological information sources, we visualized the attention scores extracted from the last attention header in the final retention layer of scMGFGRN. We investigated representative transcription factor–target gene pairs that were correctly identified as positive regulatory interactions. Two experimentally supported interactions, including KLF6-JUN and MCM7-RIF1, were selected as examples. KLF6 and JUN have been reported to participate in transcriptional regulation associated with cellular stress responses and gene expression modulation, suggesting the importance of their regulatory interaction in maintaining cellular states[45]. As shown in Fig.7E, the attention map of KLF6-JUN revealed that scRNA-seq and pseudotime features showed stronger query-side attention scores, suggesting their important roles in extracting transcriptional variations and dynamic regulatory signals. Meanwhile, GO hierarchical relationships achieved the highest aggregation score among all biological sources, indicating that functional hierarchical knowledge provides substantial guidance for accurate regulatory network inference. This finding confirms that the GO structure provides critical query information, thereby significantly enhancing model performance. In addition, the attention map of MCM7-RIF1 is visualized in Fig.7 D, it revealed distinct information exchange patterns among heterogeneous biological sources. Compared with the KLF6-JUN case, the retention affinity map of MCM7-RIF1 exhibited a different information integration pattern. For this interaction, sequence features and GO annotations showed relatively stronger contributions, highlighting the importance of GO hierarchical structure and sequence-level information in supporting the prediction of MCM7-RIF1 regulatory association.

These cases demonstrate that the proposed framework can effectively integrate heterogeneous biological information to identify reliable transcription factor–target gene interactions, providing improved interpretability for gene regulatory network reconstruction. The attention map was generated by aggregating the scores across multiple heads, providing an interpretable view of information exchange among scRNA-seq features, GO hierarchical relationships, promoter sequences, and pseudotime features during multi-source feature fusion.

### G. Potential GRN inference and validation

In this section, we applied scMGFGRN to the hESC and mHSC-E benchmark datasets from nonspecific networks (TFs + 500) to predict potential regulatory relationships. We selected three TFs with important functions and a large number of gene associations in each dataset and then selected the top 20 TF– gene pairs according to the predicting scores. As shown in Tables 2 and 3, we identified three functionally significant transcription factors and many novel TF-target relationships for both human and mouse following the same selection criteria. The inferred regulatory relationships for the hESC dataset are illustrated in Table 2. Twenty-eight pairs were successfully validated by the ground-truth dataset. For predictions not found in the ground-truth set, we consulted the hTFtarget database [39], which serves as a standardized reference for human transcription factor regulatory targets. Twelve pairs were confirmed by the hTFtarget database. An additional four pairs were corroborated through supporting evidence in literature and other functional databases. Table 3 shows the top 20 TF–gene pairs from three TFs in the mHSC-E 500 dataset. Twenty-nine pairs were directly confirmed by the ground-truth dataset. TFLink [46] is a general database that uniquely provides comprehensive and highly accurate information on mouse TF-target gene interactions. Twenty-three pairs were validated using the TFlink data-base and four pairs were further supported by specific literature. Several other predicted targets are currently classified as “unconfirmed” within these specific databases for further research.

## V. Conclusion

In this study, we developed scMGFGRN, a multi-model feature fusion deep learning framework designed to overcome the limitations of single-modality inference by integrating data-driven single-cell transcriptomic profiles with established pseudotime series, GO hierarchical relationship and gene promoter sequence. A key strength of scMGFGRN lies in its multi-modal integration strategy. Our comprehensive analysis on multiple scRNA-seq benchmarking datasets demonstrates that the inclusion of GO hierarchical relationships and semantic informatic and promoter sequence provides necessary structural information and enhances the performance of the model. In addition, the integration of GAT, BERT-based feature fusion and Gated MHA plays a critical role in the mutli-model feature extraction and lead a superior performance of scMGFGRN. The DAE effectively reduces the impact of technical noise and sparsity inherent in single-cell transcriptomic data by learning robust latent representations, thereby providing more reliable input features for downstream regulatory inference. Meanwhile, the Bi-GRU module captures dynamic regulatory patterns from pseudotime-ordered transcriptomic profiles. Unlike conventional approaches that treat single-cell expression profiles as static observations, Bi-GRU models the sequential dependencies among different cellular states along developmental trajectories, enabling scMGFGRN to characterize potential temporal regulatory changes during cell-state transitions. Furthermore, the GAT module enhances biological graph knowledge representation by explicitly modeling the structural relationships among functional annotations. Instead of treating GO terms as independent categorical features, GAT leverages the inherent graph topology of biological knowledge networks to capture hierarchical dependencies. The pretrained DNABERT-2 model further leverages gene sequence features and jointly learns associations among GRNs reconstruction.

Subsequently, the Gated MHA module adaptively assigns importance weights to different biological features, enabling the model to focus on regulatory-informative signals while suppressing irrelevant noise. This dynamic feature selection mechanism suggests that scMGFGRN does not rely on a singleinformation source but instead flexibly integrates complementary biological evidence according to specific regulatory relationships. The interpretability analysis of scMGFGRN provides further insights into the contribution of mutli-model biological information during GRN reconstruction. Attention weights results demonstrate that different regulatory interactions exhibit distinct information dependency patterns. For example, some TF-target relationships are primarily supported by transcriptomic and pseudotime features, whereas others benefit more from GO hierarchical knowledge or promoter sequence-level information. These findings indicate that scMGFGRN can adaptively identify the most informative biological signals for different regulatory contexts, improving both prediction performance and model interpretability. Further investigations demonstrate that scMGF-GRNS is capable of reconstructing cell-type-specific GRNs, predicting potential transcriptional regulatory relationships, The results consistently demonstrated that scMGF-GRN not only accurately recovers well-known regulatory interactions validated by ChIP-seq and motif databases, but also reliably predicts novel edges that are biologically plausible. Collectively, these findings exhibit the generalization capacity of scMGF-GRN, making it a robust tool for deciphering regulatory relationships in diverse biological scenarios.

Despite its promising performance, scMGFGRN still has several opportunities for future improvement. Future studies could incorporate additional regulatory modalities, such as chromatin accessibility (ATAC-seq), CITE-seq, spatial transcriptomics, and other single-cell multi-omics datasets to improve the performance of the model. scMGFGRN serves as a computational model for offering an accurate solution in GRN inference and thus contributes to decode the regulatory pattern governing cellular identity and disease progression.

## Appendix

All datasets utilized in this study are publicly accessible. The scRNA-seq and benchmark regulatory networks were obtained from the BEELINE repository [18]. The gene promoter sequence was collected from GENCODE (https://www.gencodegenes.org/). The GO hierarchy and annotation were curated from https://geneontology.org/docs/download-ontology/. The scRNA-seq dataset for human PBMC8k profiles were sourced from the 10x Genomics (https://www.10xgenomics.com/datasets/8-k-pbm-cs-from-a-healthy-donor-2-standard-2-1-0).

The ground truth regulatory networks of PBMCs and potential GRNs validation were downloaded from hTFtarget database (https://guolab.wchscu.cn/hTFtarget#!/download). The ground truth network for mouse potential GRNs validation were downloaded from TFlink (https://tflink.net/download/). The scRNA-seq data and implementation of the proposed framework scMGFGRN, including the source code for model training, data pre-processing, and GRN inference, has been made available at https://github.com/zhanglab57/scMGFGRN.git. Detailed information regarding the seven gene expression datasets and their ground-truth networks is summarized in Supplementary Table S1-2.

## A. Abbreviations and Acronyms

Abbreviation: Full name
GRNs: Gene Regulatory Networks
TF: Transcription Factors
TGI: Transcription Factor-Gene Interactions
scMGFGRN: Single-cell and Multi-model Gated feature Fusion Gene Regulatory Network Software
ANN: Artificial Neural Network
GRNBoost2: Gene Regulatory Network Boosting 2
scRNA-seq: Single-Cell RNA Sequencing
SCENIC: Single-Cell Regulatory Network Inference and Clustering
ROC: Receiver Operating Characteristic
AUROC: Area Under Receiver Operating Characteristic Curve
AUPRC: Area Under Precision-Recall Curve
BEELINE: Benchmarking Algorithms for Gene Regulatory Network Inference
ChIP-Seq: Chromatin Immunoprecipitation Sequencing
ChIP-Array: Chromatin Immunoprecipitation Array
GEO: Gene Expression Omnibus

**Lin Zhong** received master’s degree in information engineering from the Guangxi Minzu University in July 2023. He is currently working toward the PhD degree in computer science and technology with the Hunan Normal University. Currently, he is affiliated with the Department of Information Science and Engineering, Hunan Normal University. His main research interests are bioinformatics and machine learning on gene regulatory network construction and drug discovery.

**Bin Yan** is affiliated with the Faculty of Dentistry at The University of Hong Kong. He is a member of the Division of Applied Oral Sciences and Community Dental Care. His research focuses on the development and application of computational biology and bioinformatics methods for integrative omics data analysis, as well as transcriptional and epigenetic regulatory networks

**Junwen Wang** received his BS degree from Huazhong Agricultural University, his MS degrees from Jiangnan University and the University of Pennsylvania, and his PhD degree from the University of Washington. He was formerly a Professor of Biomedical Informatics at the Mayo Clinic and is currently a Professor at the Faculty of Dentistry, The University of Hong Kong.

**Minzhu Xie** received his M.S. degree in Computer Application Technology from the School of Information Science and Engineering at Central South University in 2003, and his Ph.D. degree in Computer Application Technology from the same university in 2008. He was a postdoctoral researcher at both Central South University and the University of California, Riverside. He is currently a Professor and Ph.D. Supervisor at the School of Information Science and Engineering, Hunan Normal University. His research focuses on computer algorithm design and bioinformatics

